# Subjective sleep quality in healthy young adults moderates associations of sensitivity to punishment and reward with functional connectivity of regions relevant for insomnia disorder

**DOI:** 10.1101/2025.01.31.635885

**Authors:** Michal Rafal Zareba, Tatiana Davydova, María-Ángeles Palomar-García, Jesús Adrián-Ventura, Victor Costumero, Maya Visser

**Author notes:** Correspondence (MRZ). These authors equally co-supervised the work (Victor Costumero, Maya Visser).

## Abstract

Chronic unhealthy sleeping behaviours are a major risk factor for the emergence of mood and anxiety disorders. Nevertheless, we are still lacking understanding why some individuals are more prone than others to affective dysregulation caused by sleep disruption. With preliminary evidence suggesting that brain activity during positive and negative emotional processing might play an important modulating role, we conducted whole-brain resting-state functional connectivity analyses in a large cohort of healthy young adults (N = 155). Using regions consistently affected in insomnia disorder as seeds, we investigated sleep quality-related neural connectivity patterns that were both insensitive and sensitive to the interactions with individual measures of reward and punishment processing, additionally assessing the links with indices of emotional health. The majority of the findings reflected interactions between sleep quality and reinforcement sensitivity, with the opposite associations reported in the good and poor sleepers. One of such connections, the coupling between precentral gyrus and posterior insula, was additionally negatively linked to trait anxiety, with the lowest connectivity values observed in poor sleepers with higher sensitivity to punishment. In turn, the only finding associated solely with sleep quality, i.e. coupling between subgenual anterior cingulate cortex and thalamus, was also related to the habitual use of emotion suppression strategies. As such, the present study provides evidence that reinforcement sensitivity plays an essential role in understanding the associations of poor sleep quality with brain connectivity and emotional health, hinting at a potential link that may help explain individual differences in susceptibility to sleep-related affective dysregulation.

**Highlights:**

- Functional connectivity of insomnia-related brain regions is altered in poor sleepers
- Sleep quality interacts with reinforcement sensitivity for most of such effects
- Distinct sleep-related mechanisms are linked to trait anxiety and emotion suppression
- Poor sleep and high punishment sensitivity increase the risk of affect dysregulation.

## 1. Introduction

During adolescence and early adulthood we undergo a shift towards later hours of waking up and falling asleep (Pifer et al., 2024). As the societal norms demand early hours of schooling and work, the youth are particularly susceptible to suffer from the circadian rhythm and sleep disturbances, with adverse consequences affecting them on rapid (Killgore, 2010; Owens et al., 2021; Tempesta et al., 2020), intermediate (Meyer et al., 2024; Zareba et al., 2024b) and lifespan time scales (Liu et al., 2022; Cullell et al., 2021; Hedges et al., 2024). Within hours to days, sleep deprivation causes deterioration of attention and executive functions (Killgore, 2010), leading to worse academic performance (Owens et al., 2021) on one hand, and dysregulated emotional processing on the other (Tempesta et al., 2020). Persistent inadequate sleep timing and sleep debt can eventually contribute to the emergence of mood and anxiety disorders in susceptible individuals (Meyer et al., 2024). Chronic unhealthy sleeping behaviours are further related to structural brain differences (Zareba et al., 2024b), accelerated brain ageing (Liu et al., 2022) and increased prevalence of neurodegenerative diseases (Cullell et al., 2021). With increasing amount of evidence suggesting that mental health issues experienced during youth can increase the risk of pathological brain aging (Hedges et al., 2024), understanding the mechanisms through which sleep quality is related to one’s emotional functioning can guide clinical research on interventions to improve the quality of life during both early and late stages of adulthood.

Studies have found partially convergent patterns of functional alterations in the affective brain circuitry in short-term and chronic insomnia patients (Ma et al., 2021), with some works extending such characteristics to non-diagnosed individuals experiencing sleeping difficulties (Prather et al., 2013; Baglioni et al., 2014), suggesting that processes responsible for affective dysregulation may start well before one meets diagnosis criteria. Previous functional connectivity works linking sleep-related alterations in neural circuitry with emotion regulation and mental health indices were, however, limited in their scope only to specific canonical resting-state networks (Yin et al., 2024; Chen et al., 2024; Shen et al., 2024). Importantly, the more complex mental processes, such as regulation of one’s affective state, typically require cooperation between associative areas belonging to more than one such network (Morawetz et al., 2020), which previously has not been the focus of research in the context of sleep.

Furthermore, we are still lacking understanding why some individuals are more prone than others to affective dysregulation caused by sleep disruption (Chen et al., 2017; Minkel et al., 2012; Vargas and Lopez-Duran, 2017; Wright et al., 2007). One study reported that higher levels of reward-related activity could act as a buffer against depressive symptoms associated with poor sleep (Avinun et al., 2017). Similarly, the activity of anterior insula was found to moderate the associations between negative life stress, sleep disturbances and mood symptomatology, with stronger links identified at higher levels of insular activation (Kim et al., 2022). These two studies therefore point out to a possibility that the individual sensitivity to reward and punishment may be essential for understanding the connection between poor sleep quality and affective functioning.

Therefore, the current work aims to fill this important gap in the literature, comparing for the first time sleep quality-related neural connectivity patterns, potentially within and across canonical resting-state networks, that are both insensitive and sensitive to the interactions with individual indices of reward and punishment processing. Focusing on a set of regions consistently affected in insomnia disorder (Wu et al., 2020), we conducted whole-brain resting-state functional connectivity analysis in a large cohort of healthy young adults (N = 155) with approximately half of them reporting sleeping problems. We assessed whether such neural connectivity patterns would also be related to indices of emotional well-being: the habitual use of adaptive (cognitive reappraisal) and maladaptive (suppression) emotion regulation strategies, and trait-anxiety.

Given the evidence outlined above, we expected to see differences in functional connectivity that would be dependent solely on sleep quality, as well as its interactions with indices of reinforcement sensitivity. In the case of neural patterns associated solely with poor sleep quality, we expected them to be additionally correlated with higher levels of trait-anxiety and emotion suppression, and lower levels of emotion reappraisal. Regarding the connectivity patterns linked to the interactions between sleep quality and reinforcement sensitivity, the analysis had an exploratory character as the combined effects of these measures were difficult to predict a priori due to the potential buffering effects of positive and negative emotion processing strength (Avinun et al., 2017; Kim et al., 2022).

## 2. Methods

### 2.1. Participants

The study was performed using the data of 155 healthy volunteers (22.23 ± 2.75 years, 61 males) recruited mainly from the university community via local announcements and word of mouth. All participants were right-handed as assessed by the Edinburgh Handedness Inventory (Oldfield, 1971), had normal or corrected to normal vision, no history of neurological and psychiatric disorders, and were drug-free. Self-reported sleep quality during the past month was assessed with the Pittsburgh Sleep Quality Index (PSQI; Buysse et al., 1989). Using the original criteria of PSQI scores higher than 5 indicating sleep disturbances, the sample was divided into the good (N = 86) and poor (N = 69) sleepers groups. Individual differences in the negative and positive reinforcement sensitivity were assessed using the Behavioral Inhibition and Behavioral Activation System (BIS/BAS; Carver and White, 1994), and Sensitivity to Punishment and Sensitivity to Reward Questionnaire (SPSRQ; Torrubia et al., 2001). Trait-anxiety was measured with the State-Trait Anxiety Inventory (STAI; Spielberger, 1989), while the use of reappraisal and suppression strategies for emotion regulation was tested with Emotion Regulation Questionnaire (ERQ; Gross and John, 2003; Cabello et al., 2013). All behavioural data were collected online via Qualtrics XM. The comparison of the demographic and psychological characteristics of the two groups is presented in Table 1. The poor sleepers reported higher anxiety levels and increased use of emotion suppression strategy.

**Table 1.** Demographic and psychological characteristics of the good and poor sleeper groups. Unless specified otherwise, the significance values come from independent samples t-tests. Bolded values represent significant group differences.

| Variable | Good sleepers (n = 86)<br>mean ± SD | Poor sleepers (n = 69)<br>mean ± SD | p value |
| --- | --- | --- | --- |
| Sex | 37 males, 49 females | 24 males, 45 females | 0.297 <sup>a</sup> |
| Age | 22.43 (2.61) | 21.97 (2.91) | 0.302 |
| PSQI | 3.70 (1.15) | 8.58 (2.65) | **< 0.001** |
| BIS | 21.62 (3.33) | 21.96 (3.42) | 0.533 |
| BAS | 41.62 (5.03) | 40.91 (5.16) | 0.394 |
| SP | 10.90 (5.36) | 11.87 (5.20) | 0.257 |
| SR | 9.56 (3.91) | 10.16 (3.97) | 0.346 |
| STAI | 21.16 (9.37) | 26.54 (12.22) | **0.002** |
| ERQ reappraisal | 29.94 (6.14) | 28.36 (6.49) | 0.123 |
| ERQ suppression | 13.74 (5.81) | 16.17 (5.76) | **0.01** |

<sup>a</sup> Chi-squared test.
Abbreviations: SD, standard deviation; PSQI, Pittsburgh Sleep Quality Index; BIS, Behavioral Inhibition System; BAS, Behavioural Activation System; SP, sensitivity to punishment; SR, sensitivity to reward; STAI, State-Trait Anxiety Inventory; ERQ, Emotion Regulation Questionnaire.

### 2.2. MRI data acquisition and preprocessing

Two types of images were collected for each participant using a 3 Tesla scanner (SIGNA Architect, GE Healthcare, Chicago, Illinois, United States) and a 24-channel head coil. High-resolution anatomical image was obtained with T1 BRAVO sequence (384 sagittal slices; 0.5 × 0.47 × 0.47 mm voxel size; TR = 8.516 ms; TE = 3.276 ms). The resting-state fMRI data was collected with a T2*-weighted gradient echo planar imaging sequence (27 axial slices acquired in the interleaved bottom-up order; 3.75 × 3.75 × 4.5 mm voxel size; TR = 2000 ms; TE = 30 ms; flip angle = 70°; 210 volumes resulting in 7 mins of data). During the functional scan, the participants were instructed to keep their eyes open and concentrate on the fixation cross without thinking about anything in particular.

Preprocessing of the data was performed using CONN release 21.a (Whitfield-Gabrieli and Nieto-Castanon, 2012) and SPM release 12.7771 (Penny et al., 2011). The modular preprocessing pipeline (Nieto-Castanon, 2020) included realignment with correction of susceptibility distortion interactions, slice timing correction, outlier detection, direct segmentation and MNI-space normalisation, and smoothing. Functional data were realigned using SPM realign and unwarp procedure (Andersson et al., 2001), where all scans were coregistered to a reference image (first scan of the first session) using a least squares approach and a 6 parameter (rigid body) transformation (Friston et al., 1995), and resampled using b-spline interpolation to correct for motion and magnetic susceptibility interactions. Temporal misalignment between different slices of the functional data was corrected following SPM slice-timing correction procedure (Sladky et al., 2011), using sinc temporal interpolation to resample each slice BOLD time series to a common mid-acquisition time. Potential outlier scans were identified using ART (Whitfield-Gabrieli et al., 2011) as acquisitions with framewise displacement above 0.9 mm or global BOLD signal changes above 5 standard deviations (Power et al., 2014), and a reference BOLD image was computed for each subject by averaging all scans excluding outliers. Functional and anatomical data were normalised into standard MNI space, segmented into grey matter (GM), white matter, and cerebrospinal fluid (CSF) tissue classes, and resampled to 3 mm isotropic voxels following a direct normalisation procedure (Calhoun et al., 2017) using SPM unified segmentation and normalisation algorithm (Ashburner and Friston, 2005; Ashburner, 2007) with the default IXI-549 tissue probability map template. Subsequently, functional data were smoothed using spatial convolution with a Gaussian kernel of 4 mm full width half maximum (FWHM). Lastly, functional data were denoised using a standard denoising pipeline including the regression of potential confounding effects characterised by white matter timeseries (5 CompCor noise components), CSF timeseries (5 CompCor noise components), motion parameters and their first order derivatives (12 factors; Friston et al., 1996), outlier scans (below 52 factors; Power et al., 2014), and linear trends (2 factors), followed by bandpass frequency filtering of the BOLD time series (Hallquist et al., 2013) between 0.008 Hz and 0.09 Hz. CompCor (Behzadi et al., 2007; Chai et al., 2012) noise components within white matter and CSF were estimated by computing the average BOLD signal as well as the largest principal components orthogonal to the BOLD average, motion parameters, and outlier scans within each subject’s eroded segmentation masks. From the number of noise terms included in this denoising strategy, the effective degrees of freedom of the BOLD signal after denoising were estimated to range from 40.7 to 57.7 (average 57.2) across all subjects.

### 2.3. Regions of interest and seed-based connectivity analysis

Regions of interest (ROIs) for the seed-based functional connectivity analyses were selected based on the results of the recent meta-analyses on the structural and functional brain changes in insomnia disorder (Wu et al., 2020). ROIs were defined as 4 mm-radius spheres centred on the peak coordinates of reported differences in GM volume, glucose metabolism, amplitude of low frequency fluctuations (ALFF) and regional homogeneity (ReHo). The only exception from this rule was made for anterior cingulate cortex (ACC) as the meta-analysis results partially mapped onto the border between subgenual (sgACC) and pregenual ACC (pgACC), which are known to engaged more in the processing of, respectively, positive and negative reinforcement (Rolls, 2019). In this case, we decided to use larger, atlas-based ROIs, distinguishing between the ACC subregions (Fan et al., 2016). The ROIs with their mass centre coordinates are presented in Table 2. If more than one ROI was present in a given anatomical location in one hemisphere, the corresponding Brodmann areas (BA) were added for easier discrimination.

**Table 2.**
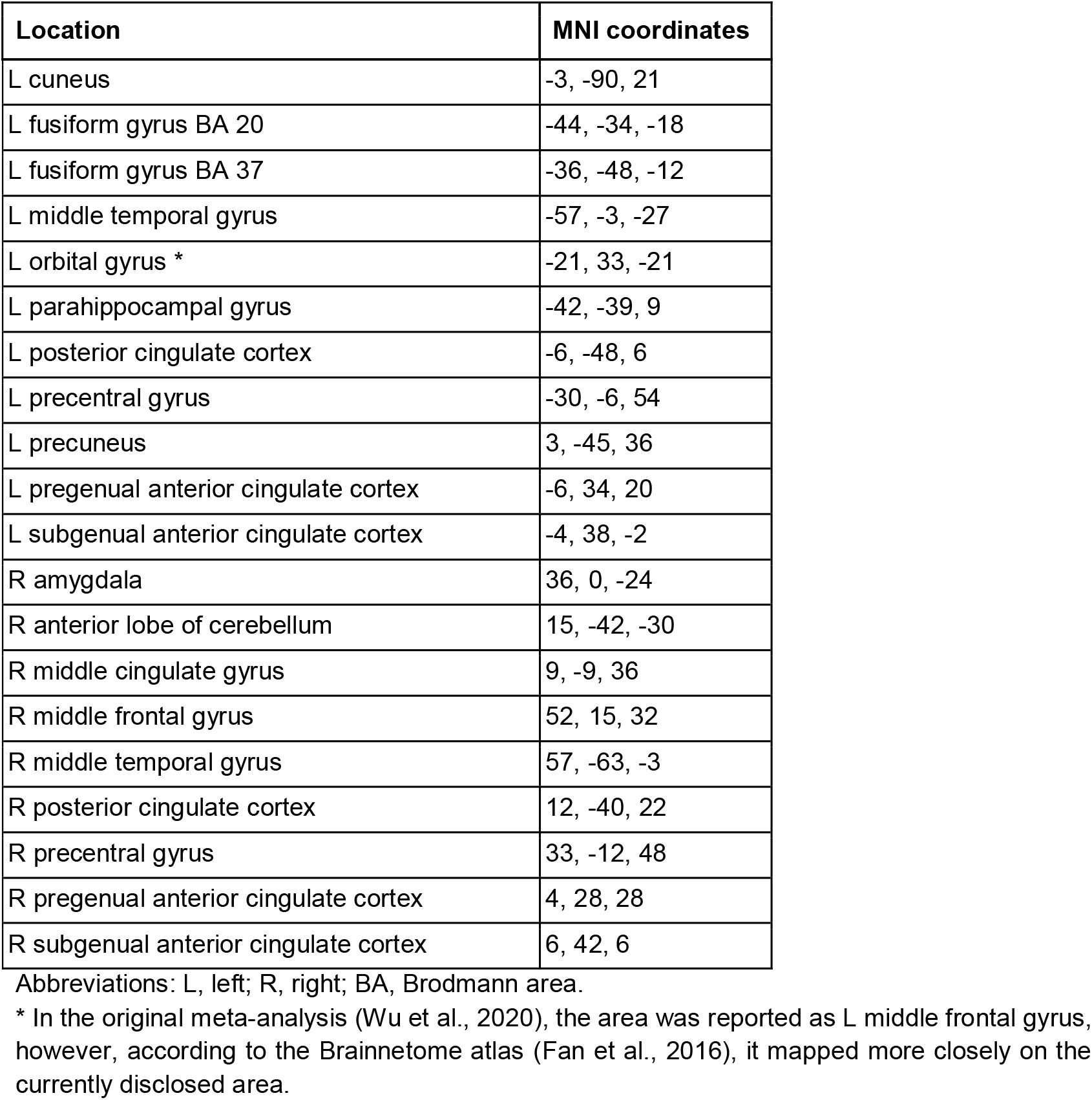
Regions of interest (ROIs) for the seed-based functional connectivity analyses.

| Location | MNI coordinates |
| --- | --- |
| L cuneus | -3, -90, 21 |
| L fusiform gyrus BA 20 | -44, -34, -18 |
| L fusiform gyrus BA 37 | -36, -48, -12 |
| L middle temporal gyrus | -57, -3, -27 |
| L orbital gyrus * | -21, 33, -21 |
| L parahippocampal gyrus | -42, -39, 9 |
| L posterior cingulate cortex | -6, -48, 6 |
| L precentral gyrus | -30, -6, 54 |
| L precuneus | 3, -45, 36 |
| L pregenual anterior cingulate cortex | -6, 34, 20 |
| L subgenual anterior cingulate cortex | -4, 38, -2 |
| R amygdala | 36, 0, -24 |
| R anterior lobe of cerebellum | 15, -42, -30 |
| R middle cingulate gyrus | 9, -9, 36 |
| R middle frontal gyrus | 52, 15, 32 |
| R middle temporal gyrus | 57, -63, -3 |
| R posterior cingulate cortex | 12, -40, 22 |
| R precentral gyrus | 33, -12, 48 |
| R pregenual anterior cingulate cortex | 4, 28, 28 |
| R subgenual anterior cingulate cortex | 6, 42, 6 |

Abbreviations: L, left; R, right; BA, Brodmann area.
\* In the original meta-analysis (Wu et al., 2020), the area was reported as L middle frontal gyrus, however, according to the Brainnetome atlas (Fan et al., 2016), it mapped more closely on the currently disclosed area.

Seed-based connectivity analyses were performed in AFNI (Cox, 1996). For every ROI in each participant, we extracted the mean blood-oxygen-level-dependent (BOLD) signal time series, and computed the Pearson correlations with the time courses of all other brain voxels. The distribution of the correlation coefficients was normalised with the Fisher’s z transform. Whole-brain functional connectivity maps created in this manner were used as the input to the statistical analyses.

### 2.4. Statistical analysis

All neuroimaging analyses were performed with the 3dMVM (Chen et al., 2014) package available as a part of AFNI (Cox, 1996). For each ROI, we tested an ANCOVA-style design in which its functional connectivity was predicted by the subjective sleep quality (good or poor sleeper group), sensitivity to reward (SR) and sensitivity to punishment (SP), modelling the main effects, as well as the two- and three-factor interactions. Biological sex and age of participants were controlled as covariates. The Results section will include all the main effects, two-term and three-term interactions that survived multiple comparison correction on the cluster-level, which was achieved through family-wise error rate correction (FWE < 0.05) after the initial voxel-level thresholding at the level of p < 0.001 (Flandin and Friston, 2019). The cluster size threshold for each analysis was determined by first using 3dFWHMx’s spatial autocorrelation function to estimate the smoothness of the noise in the residuals files, and subsequently providing these parameters as the input into 3dClustSim.

To determine the nature of the significant interactions from the aforementioned analyses, post-hoc tests were applied in R (version 4.2.1; R Core Team, 2022). For the two-way interactions, slope differences between poor and good sleepers groups were calculated with the *emtrends* function from the *emmeans* library. For the three-way interactions, simple slopes analysis was run with the *sim_slopes* function from the *interactions* package to compare slopes at different values of SP across the two groups.

To test the replicability of the reported effects across the psychometric scales, this time using BIS/BAS (Carver and White, 1994), we extracted the connectivity patterns successfully predicted in the whole-brain models and used them as the dependent variables in permutation-based ANCOVAs (10000 permutations). The decision to use a non-parametric approach was guided by the lack of normal distribution of the BIS data. The calculations were run using the *aovperm* function from the *permuco* library (Frossard and Renaud, 2021) in R, including biological sex and age as covariates, akin to the original analyses. For illustrative purposes only, the magnitude of the effects across the main and replication analyses was determined using partial eta-squared (ηp2). It should be noted that the magnitudes of the effects reported in the whole-brain analyses are likely to be overestimated, which is an inherent limitation of the mass univariate fMRI analysis (Kriegeskorte et al., 2010).

We additionally tested whether the highlighted connectivity patterns were associated with trait anxiety and the use of emotion reappraisal and suppression strategies. This analysis was meant to elucidate whether the same neural patterns that were associated with sleep quality and its interactions with reinforcement sensitivity measures were also related to negative emotionality and maladaptive emotion regulation. Multiple comparison correction across the replication and associative analyses were achieved with the false discovery rate (FDR < 0.05) approach.

## 3. Results

### 3.1. Effects of sleep quality and reinforcement sensitivity on neural connectivity

The whole-brain analyses yielded five significant findings, one for the main effect of subjective sleep quality and four others for the interactions with SP and SR (Table 3). Poor sleepers were found to have lower functional connectivity between the right sgACC and left prefrontal thalamus (Figure 1A). In turn, interactions between the sleep quality and SP predicted the connectivity of the left precentral gyrus with posterior insula (Figure 2A), and supramarginal and superior temporal gyri of the same hemisphere (Figure 3B). Similarly, the functional connectivity of the left fusiform gyrus BA 20 and bilateral supplementary motor area was dependent on the interaction between the sleep quality and SR (Figure 3A). In all the cases, the reinforcement sensitivity measures were positively linked to functional connectivity in good sleepers, while the opposite relationships were observed for poor sleepers. The post-hoc tests confirmed significant slope differences between the groups for all the connectivity patterns associated with the two-way interactions (Supplementary Table 1).

**Table 3.** Results of the seed-based functional connectivity analyses. Multiplication sign (×) indicates interactions between the factors.

| Seed | Factors | MNI | Location | Voxels | F-stat | $\eta p^2$ |
| --- | --- | --- | --- | --- | --- | --- |
| R subgenual anterior cingulate cortex | Sleep quality | -12, -16, 8 | L prefrontal thalamus | 37 | 19.66 | 0.143 |
| L fusiform gyrus BA 20 | Sleep quality × SR | 10, 8, 56 | B supplementary motor area | 46 | 21.59 | 0.154 |
| L precentral gyrus | Sleep quality × SP | -56, -22, 14 | L supramarginal gyrus<br>L superior temporal gyrus | 80 | 24.86 | 0.164 |
| L precentral gyrus | Sleep quality × SP | -42, -10, 0 | L posterior insula | 69 | 31.64 | 0.187 |
| R middle temporal gyrus | Sleep quality × SP × SR | -32, 50, 12 | L lateral prefrontal cortex | 86 | 23.99 | 0.173 |

Abbreviations: $\eta p^2$ , partial eta-squared; L, left; R, right; B, bilateral; BA, Brodmann area; SR, sensitivity to reward; SP, sensitivity to punishment.

**Figure 1.**
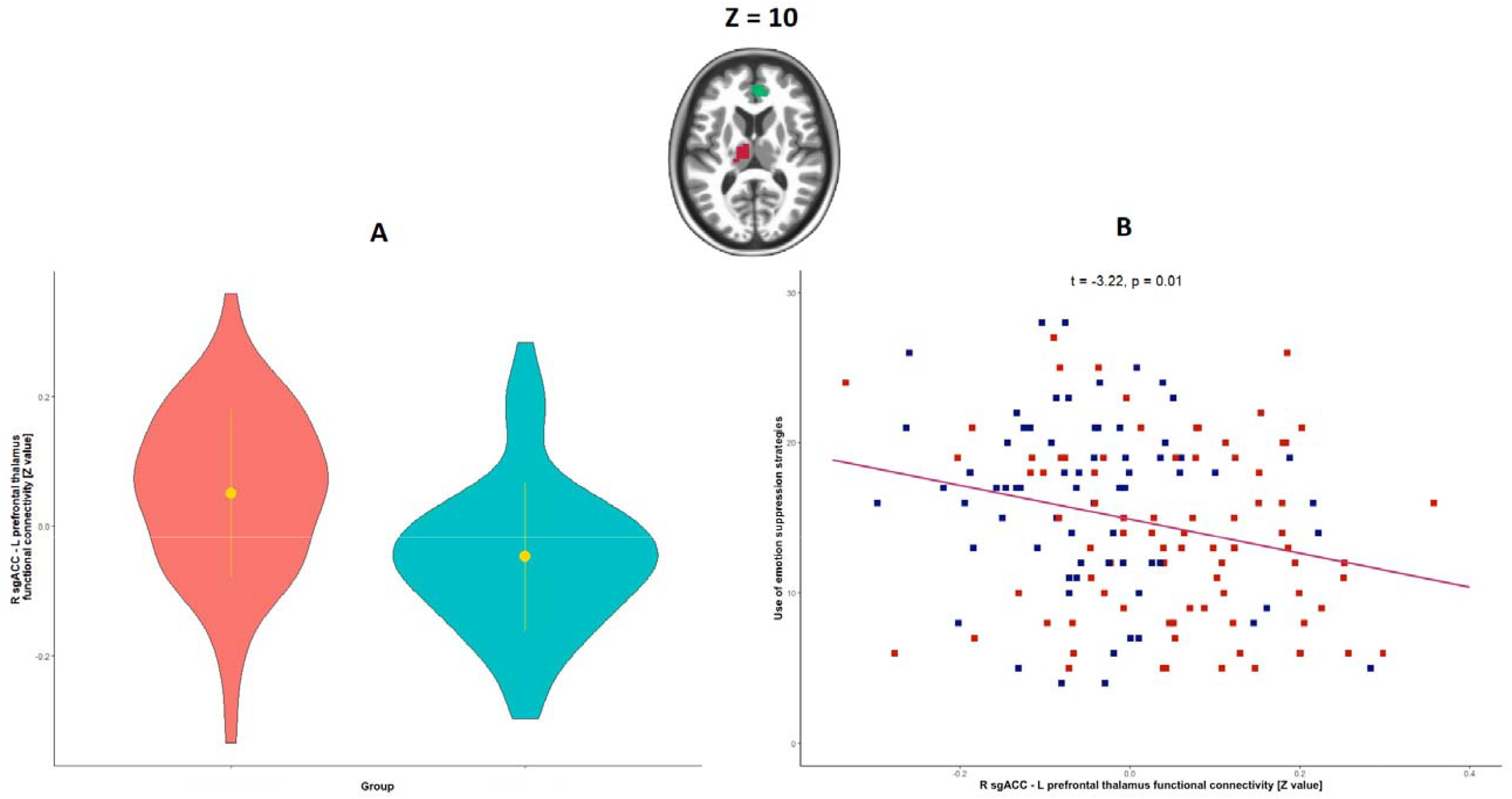
(A) Poor sleepers (blue) were found to have lower functional connectivity between the right subgenual anterior cingulate cortex (R sgACC) and the left prefrontal thalamus compared to the good sleepers (red). (B) Functional connectivity between the two areas was additionally negatively associated with the habitual use of emotion suppression strategies. For illustrative purposes, we used the average of connectivity values from the voxels within the thalamus cluster. R sgACC seed is presented in green, while the left prefrontal thalamus cluster is shown in magenta.

**Figure 2.**
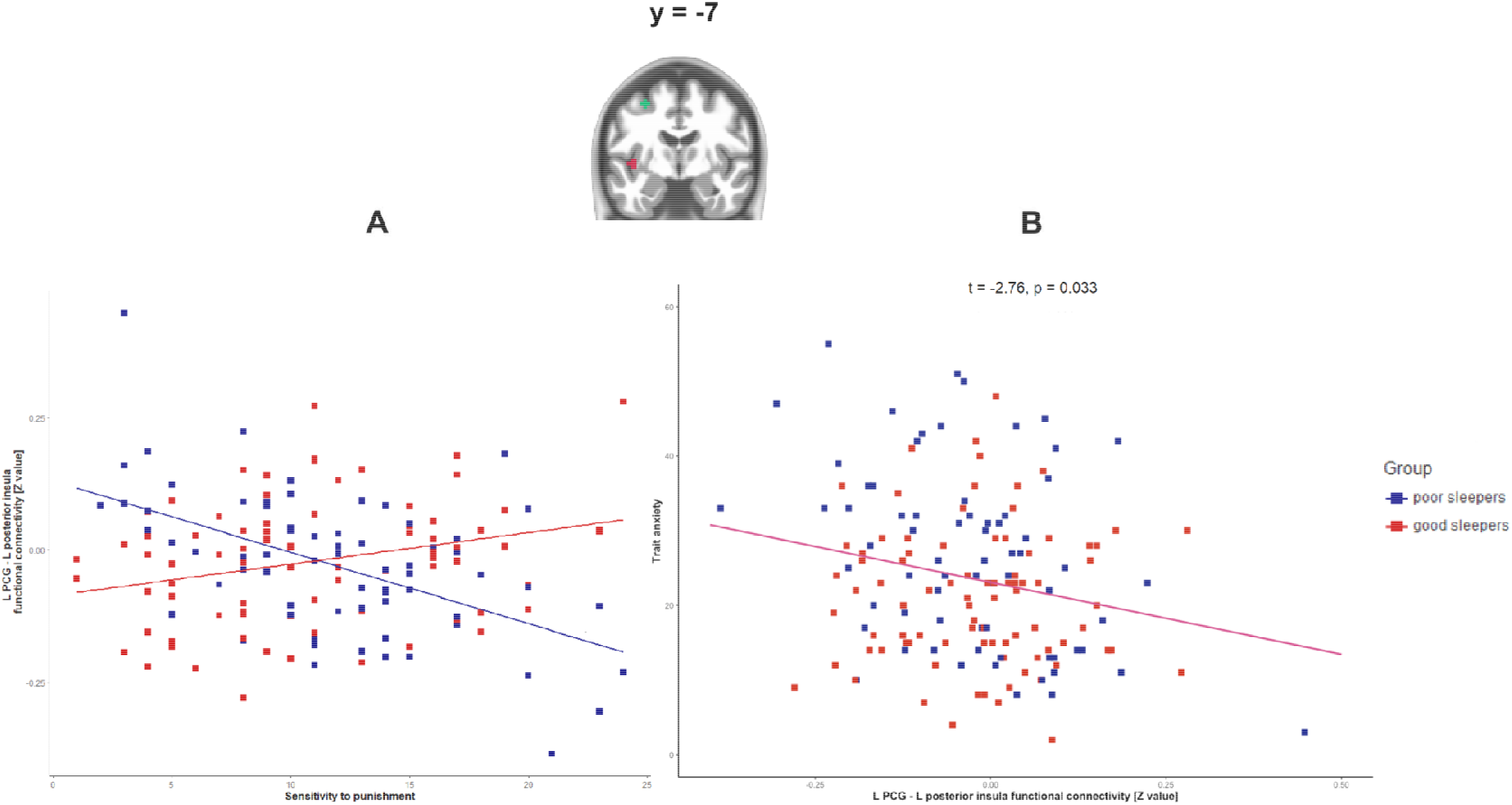
The functional connectivity of the left precentral gyrus (L PCG) with the left posterior insula was sensitive to the interactions of individual sleep quality with negative reinforcement sensitivity (A). The strength of the coupling between the areas was additionally negatively correlaed with trait anxiety (B). For illustrative purposes, we used the average of connectivity values from the voxels within the insula cluster. L PCG seed is presented in green, while the left posterior insula cluster is shown in magenta.

**Figure 3.**
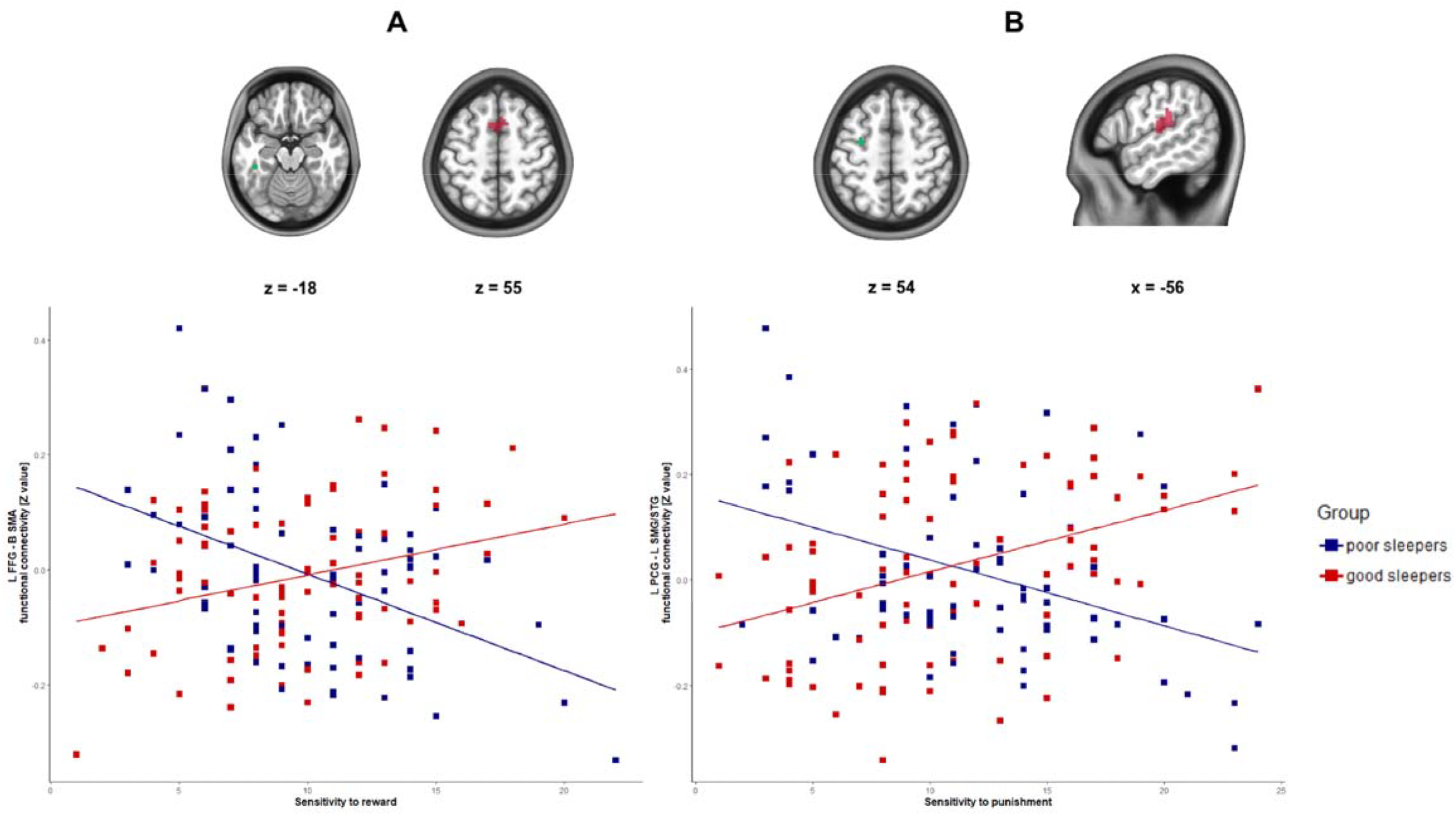
The associations between reinforcement sensitivity measures and functional connectivity patterns were moderated by individual sleep quality. Significant findings for the positive reinforcement (A) were observed for the connectivity between the left fusiform gyrus (L FFG) and bilateral supplementary motor area (B SMA). As for the negative reinforcement (B), such associations were found for the connectivity of the left precentral gyrus (L PCG) with the left supramarginal and superior temporal gyri (L SMG/STG). For illustrative purposes, we used the average of connectivity values from the significant voxels in each cluster. Seeds are presented in green, and the areas highlighted by the whole-brain analyses are shown in magenta.

In turn, a three-way interaction between sleep quality, SP and SR was observed for the functional connectivity of the right middle temporal gyrus and the left lateral prefrontal cortex (Figure 4). The post-hoc test revealed that at lower levels of SP (i.e. a standard deviation below the mean), poor sleepers had a positive relationship between SR and functional coupling between the two areas (t = 3.46; p = 0.0007). No association between the two metrics was observed at the mean level of SP (t = 0.41; p = 0.680), and a significant negative link was found at higher levels of SP (i.e. a standard deviation above the mean; t = -2.56; p = 0.012). The directionality of the relationships was reversed in the group without sleeping problems (Figure 4). At lower levels of SP, good sleepers had a negative association between SR and functional connectivity (t = -3.82; p = 0.0002). No significant links were found at either mean (t = -1.10; p = 0.271) or higher levels of SP (t = 1.65; p = 0.101) in this group.

**Figure 4.**
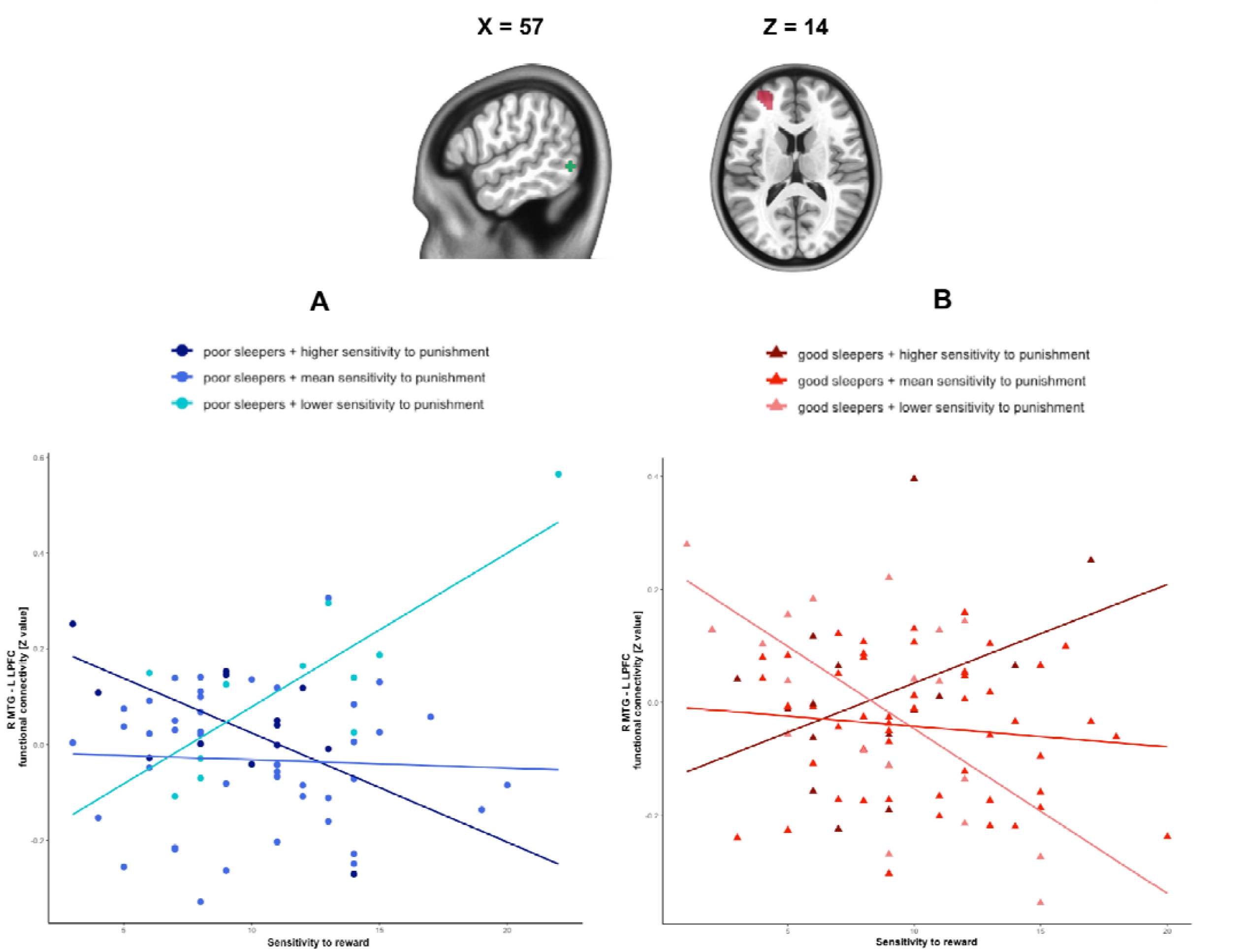
Three-way interactions between sleep quality, positive and negative reinforcement sensitivity were found for the functional connectivity between the right middle temporal gyrus (R MTG) and the left lateral prefrontal cortex (L LPFC). (A) Poor sleepers with lower sensitivity to punishment had a positive association between sensitivity to reward and the connectivity, whereas in poor sleepers with higher sensitivity to punishment the relationship was negative. (B) The direction of the associations was reversed in the good sleepers. For illustrative purposes, the good and poor sleepers groups were divided into three subgroups with higher (i.e. more than one standard deviation (SD) above the mean), lower (i.e. more than one SD below the mean) and mean sensitivity to punishment (i.e. values in between the previously described categories). Plots were drawn using average connectivity values from the voxels in the L LPFC cluster. R MTG seed is shown in green, and the frontal area in magenta.

The analysis performed with the BIS/BAS data replicated the effects of interactions between sleep quality and SR on the functional connectivity between the left fusiform gyrus BA 20 and bilateral supplementary motor area (F = 7.53; p_FDR_ = 0.03; ηp^2^ = 0.049; Supplementary Figure 1). As for the main effects of SR and SP, they were observed for 5 functional connectivity patterns each, and were characterised by higher replicability (80%) than the interaction effects. As these associations were not the primary focus of the paper, they are provided in the Supplementary Material (Supplementary Table 2 and Supplementary Figure 2) for the sake of transparency and to facilitate their reusability for meta-analytic purposes.

### 3.2. Associations of the highlighted connectivity patterns with anxiety and emotion regulation

Two of the highlighted functional connectivity patterns were also related to indices of emotional health. The strength of the coupling between the right sgACC and left prefrontal thalamus was negatively related to the use of emotion suppression strategies (t = -3.22; p_FDR_ = 0.01; Figure 1B), while the functional connectivity between the left precentral gyrus and posterior insula was negatively associated with trait anxiety (t = -2.76; p_FDR_ = 0.033; Figure 2B). All the other associations were not significant (Supplementary Table 3). The links with anxiety and emotion regulation for the connectivity patterns related to the main effects of positive and negative reinforcement measures are provided in Supplementary Table 4.

## 4. Discussion

In line with the hypotheses, the current work highlights a fact that experiencing poor sleep quality over a period of a month in a sample of healthy young adults is related to differential functional connectivity of brain areas implicated in insomnia disorder. In more detail, poor sleep quality was negatively related to the connectivity between sgACC and thalamus, with a similar association observed for the use of emotion suppression strategies. The majority of the findings, however, revealed interactions between sleep quality and reinforcement sensitivity, with opposite associations reported in good versus poor sleepers. For two-way interactions, functional connectivity patterns of the left precentral and fusiform gyri were positively associated with reinforcement sensitivity in good sleepers, whereas a negative link was observed in individuals reporting sleeping problems. In addition, the coupling between the left precentral gyrus and posterior insula was negatively linked to trait anxiety. For three-way interactions, a more complex pattern emerged in the functional connectivity of the right middle temporal gyrus with the left lateral prefrontal cortex. Poor sleepers with lower SP showed increased functional connectivity at higher SR levels, whereas in poor sleepers with higher SP the association was negative. The directionality of these links was reversed in the good sleepers.

The results of the current study are in line with previous research showing that chronic poor sleep quality is related to differential functional connectivity of brain areas within and between commonly identified networks (Chen et al., 2024; Shen et al., 2024; Zhang et al., 2020), suggesting that sleep is linked to brain function through multiple independent pathways. The prevalence of the findings associated with individual reinforcement sensitivity, as compared to a single connectivity pattern differing between the groups based solely on sleep quality, speaks to the crucial role played by the former in understanding the relationship between poor sleep and one’s well-being. Importantly, both types of connectivity patterns are also linked to separate indices of emotional health, i.e. emotion suppression and trait anxiety, respectively.

The literature suggests multidirectional causal relationships between sleep quality, emotion suppression and anxiety (Dawel et al., 2021; Vazsonyi et al., 2022; Tavernier and Willoughby, 2015; Bendal et al., 2024). For example, suppressing negative emotions prevents their conscious processing and over time can lead to chronic stress and anxiety. On the other hand, individuals with high levels of anxiety may suppress emotions to avoid emotional discomfort. While suppression can create short-term relief, reinforcing the behavior, in the long run it prevents emotional regulation, further increasing anxiety. Both processes may also contribute to poor sleep quality through increasing physiological arousal (Tyra et al., 2024; Yin et al., 2024). In turn, deteriorated sleep quality can similarly affect emotional reactivity and regulation (Cox & Olatunji, 2020). Taking into account this complex picture, by identifying separate sleep-related neural pathways related to emotion suppression and trait anxiety, our work makes a step forward towards the delineation of the circuitry responsible for each of these often overlapping psychological phenomena, with the reported functional coupling believed to reflect both the history of prior co-activation between areas but also the propensity to cooperate during future execution of cognitive processes (Guerra-Carrillo et al., 2014).

As for sgACC, mounting evidence pinpoints its pivotal role in the emotional dysregulation observed across different levels of sleep debt and in clinical populations (Motomura et al., 2013; Chai et al., 2023; Reimann et al., 2023). Given its role as an interface between emotional and cognitive processes (Reiman et al., 2023), the involvement of this area in emotional suppression comes as no surprise (Motomura et al., 2013). Interestingly, the results of the current study suggest that different sgACC-related pathways might be responsible for this phenomenon at different stages of emotion processing. For instance, accumulating sleep debt over 5 consecutive days was found to increase reactivity of amygdala to the emotional faces, a finding that was proportional to the decrease in amygdala’s connectivity to sgACC (Motomura et al., 2013). Similarly, enhanced connectivity between these two areas after a night of total sleep deprivation was associated with better mood in healthy participants and antidepressant effects in depressed patients (Chai et al., 2023). Therefore, decreased connectivity between sgACC and amygdala appears related to an increase of the latter’s activity. As the thalamic region identified in the current work is involved in sensorimotor functions (Fan et al., 2016), its lower connectivity with sgACC might reflect decreased propensity to act as a result of experienced emotions, providing a mechanism that could potentially partially counterbalance the increased amygdala reactivity (Goldin et al., 2008; Palmer and Alfano, 2017).

The remaining connectivity patterns identified in the study, involving sensory cortices and executive control areas, were associated with interactions of sleep quality and individual reinforcement sensitivity. Previous neuroimaging studies have shown that the anatomy of the sensory cortices mediated associations of stress and negative emotionality with worsened sleep quality (Zhang et al., 2022; Wang et al., 2021), additionally linking it with circadian phenotypes at risk of experiencing sleep problems (Zareba et al., 2022, Vulser et al., 2023). A recent meta-analysis has demonstrated that short-term sleep restriction (i.e. up to a week) has a modest dampening effect on emotional arousal regardless of its valence, and a large diminishing and augmenting impact on positive and negative mood, respectively (Tomaso et al., 2021). However, individual functional neuroimaging studies seem to challenge such a uniform view. Short-term sleep disruption has been reported to increase the mesolimbic system’s reactivity to pleasure-evoking stimuli, as well as anticipation and reception of financial gains (Gujar et al., 2011; Venkatraman et al., 2007; Venkatraman et al., 2011), additionally linking it to greater risk-taking (Telzer et al., 2013). On the other hand, no such associations were found in a large sample of adults with chronic poor sleep quality (Curtis et al., 2019), while in youth with anxiety sleep disturbances were related to decreased mesolimbic reactivity to monetary rewards (Sollenberger et al., 2023). Similarly, reports have indicated sleep deprivation-related increases in brain reactivity to negative emotional stimuli (Yoo et. al, 2007; Motomura et al., 2013) but also less disappointment with monetary losses (Venkatraman et al., 2011) and an impaired ability to learn from negative outcomes (Venkatraman et al., 2007). There is thus a substantial variability in the effects of sleep loss on reinforcement processing, with the duration of experiencing sleeping problems, individual differences in sensitivity to specific types of rewards and punishment and genetic variation in the dopaminergic genes being likely contributors (Tomaso et al., 2021; Greer et al., 2016).

The current work provides an insight into the potential neural basis of such a broad range of associations. The left fusiform gyrus and right middle temporal gyrus are higher-order visual cortices participating in processing words, objects and faces, and as such they contribute to the recognition of external stimuli and their meaning (Fan et al., 2016; Yarkoni et al., 2011). In turn, the bilateral supplementary motor area and the left lateral prefrontal cortex are related to various flavours of executive control functions, including emotion regulation, yet through differential involvement across a number of cognitive processes (Fan et al., 2016; Morawetz et al., 2020).

Given their roles, the functional coupling between these two types of brain regions may reflect the frontal cortex’s ability to exert control over the semantic and emotional mental representations evoked by visual cognition. With increased risk-taking in poor sleepers previously reported to be paralleled by reduced functional coupling between executive control hubs and affective circuitry (Telzer et al., 2013), the current results suggest that such an imbalance in the case of the connectivity between the left fusiform gyrus and bilateral supplementary motor area may be related to preservation of positive reinforcement sensitivity in individuals experiencing chronic sleep problems.

As for the connectivity of the left precentral gyrus with ipsilateral posterior insula, supramarginal and superior temporal gyri, all the areas belong to the pericentral sensorimotor network (Uddin et al., 2019). The involvement of these regions in negative reinforcement sensitivity is likely related to their shared role in pain processing (Yarkoni et al., 2011). Sleep deprivation has been previously linked to increased subjective pain ratings with a concurrent decrease of neural reactivity to painful stimuli (Tiede et al., 2010), a pattern mirroring our functional connectivity findings (i.e., lower coupling associated with higher SP). Interestingly, a recent meta-analysis has reported decreased pain threshold in anxious individuals (Scaini et al., 2025), indicating that the associated neural mechanisms could play a role in linking these phenomena. Indeed, one of the identified connectivity patterns, the coupling of the precentral gyrus with the posterior insula was negatively associated with trait anxiety, identifying individuals with chronic sleep problems and co-occurring high SP as a group at a particularly high risk of developing mental health issues.

Previous studies have highlighted the particular role of insula in emotion processing circuitry across subclinical and clinical anxiety (Zareba et al, 2024a; Wang et al., 2019; Pico-Perez et al., 2017). Higher trait anxiety has also been linked to greater increase in the reactivity of anterior insula to emotional cues following sleep deprivation (Goldstein et al., 2013). Therefore, it is tempting to propose that psychological interventions altering the functioning of the insula might be an effective preventive measure for individuals at increased risk of chronic sleep disruption, for example due to shift work or inflexible school timings. Indeed, the activity of this region has recently been found to be predictive of the outcome of cognitive-behavioural therapy in anxiety-related disorders (Pico-Perez et al., 2023). Nevertheless, in the context of the current cross-sectional study, we cannot disentangle whether individual affective functioning and emotion regulation changed as a result of poor sleep quality, or whether the baseline emotional traits affected the sleep perception and related alterations in functional connectivity.

Additionally, it needs to be noted that subjective poor sleep quality measured in the current study might reflect individual negative beliefs about sleep rather than actual sleep disturbances, which can in turn impact one’s affective cognition, as evidenced by the fact that challenging the negative beliefs about sleep constitutes a core part of the cognitive-behavioural therapy for insomnia. These limitations preclude drawing any conclusion on causal relationships in the current study, posing a potential limitation to the clinical usability of the findings. Future longitudinal works should address this issue by comparing how using subjective and objective measures of sleep quality in the same set of individuals, accounting for their sleep need, affects the associations of sleep and reinforcement sensitivity with brain connectivity. This is especially important given that previous literature suggests discrepancies in the neural correlates between objective and subjective sleep measures (Grumbach et al., 2020). Despite the limitations discussed above, subjective rating of sleep quality are a valuable and useful research tool as due to interindividual variability in physiological factors, such as the dissipation of homeostatic sleep pressure, quantitatively similar sleep might lead to completely different levels of qualitative subjective vigourness (Mongrain et al., 2006), making subjective measures conceptually closer related to affective functioning.

On a more methodological note, given the innovative and partially exploratory character of the study, no additional correction for the number of tested seeds was applied. As ROI-level analyses also mitigate effect size inflation (Kriegeskorte et al., 2010), future studies could address this limitation by focusing on connectivity patterns between specific, literature-based sets of regions, for example drawn from the current work. Additionally, while males and females did not differ in terms of sleep quality and emotional traits in the current sample, previous literature has suggested the existence of between-sex differences in these measures (Madrid-Valero et al., 2023; Lischke et al., 2020). Future, large-scale investigation could thus probe this idea by investigating potential moderating effects of sex on neural correlates of affective processes in the context of differential sleep quality. Last but not least, it needs to be stressed that the replicability of connectivity effects associated with interactions between sleep quality and reinforcement sensitivity was lower than for the latter’s main effects. One potential explanation for such a finding might be the difference in how single items are scored in the used questionnaires. While SPSRQ (Torrubia et al., 2001) has a binary scoring system, the approach deployed in BIS/BAS (Carver and White, 1994) permits a broader range of responses (from 1 to 4). As such, while the total scores in both measures are highly correlated with each other, the more nuanced interactions might be more difficult to observe across the two scales.

## 5. Conclusions

With the majority of the findings involving interactions with individual differences in positive and negative reinforcement sensitivity, the present study contributes to the existing literature by providing evidence that reinforcement sensitivity plays an essential role in understanding the associations of poor sleep quality with brain connectivity and emotional health. Importantly, the discussed connectivity patterns are also separately linked to trait anxiety and the use of emotion suppression strategies, indicating that different emotional facets associated with worsened sleep quality may have distinct underlying mechanisms. This is particularly underlined by the modulating effect of sensitivity to punishment on the connection between the precentral gyrus and insula, with its coupling strength additionally associated with trait anxiety. As such, the current work increases our understanding of the links between sleep and mental health, hinting at a potential mechanism that may help explain individual differences in susceptibility to sleep-related affective dysregulation (Chen et al., 2017; Minkel et al., 2012; Vargas and Lopez-Duran, 2017; Wright et al., 2007).

## Data availability

Binary masks showing the areas significantly associated with each contrast will be made available on NeuroVault after the manuscript acceptance.

## CRediT author statement

MRZ: Conceptualization, Formal analysis, Writing - Original Draft, Visualisation. TD: Investigation, Writing - Review & Editing. MAPG: Investigation, Writing - Review & Editing. JAV: Investigation, Writing - Review & Editing. VC: Conceptualisation, Formal analysis, Supervision, Writing - Review & Editing. MV: Conceptualisation, Supervision, Writing - Review & Editing.

## Ethical statement

All the procedures followed were in accordance with the ethical standards of the responsible committee on human experimentation (institutional and national) and with the Declaration of Helsinki (1975), and the applicable revisions at the time that this research was underway. Informed consent to be included in the study was obtained from all the participants. The study protocol was approved by the Universitat Jaume I Ethics Committee.

## References

Andersson, J. L., Hutton, C., Ashburner, J., Turner, R., & Friston, K. (2001). Modeling geometric deformations in EPI time series. NeuroImage, 13(5), 903–919. 10.1006/nimg.2001.0746

Ashburner J. (2007). A fast diffeomorphic image registration algorithm. NeuroImage, 38(1), 95–113. 10.1016/j.neuroimage.2007.07.007

Ashburner, J., & Friston, K. J. (2005). Unified segmentation. NeuroImage, 26(3), 839–851. 10.1016/j.neuroimage.2005.02.018

Avinun, R., Nevo, A., Knodt, A. R., Elliott, M. L., Radtke, S. R., Brigidi, B. D., & Hariri, A. R. (2017). Reward-Related Ventral Striatum Activity Buffers against the Experience of Depressive Symptoms Associated with Sleep Disturbances. The Journal of neuroscience : the official journal of the Society for Neuroscience, 37(40), 9724–9729. 10.1523/JNEUROSCI.1734-17.2017

Baglioni, C., Spiegelhalder, K., Regen, W., Feige, B., Nissen, C., Lombardo, C., Violani, C., Hennig, J., & Riemann, D. (2014). Insomnia disorder is associated with increased amygdala reactivity to insomnia-related stimuli. Sleep, 37(12), 1907–1917. 10.5665/sleep.4240

Behzadi, Y., Restom, K., Liau, J., & Liu, T. T. (2007). A component based noise correction method (CompCor) for BOLD and perfusion based fMRI. NeuroImage, 37(1), 90–101. 10.1016/j.neuroimage.2007.04.042

Bendall, R. C. A., Elton, S. N., & Hughes, A. T. L. (2024). Expressive suppression mediates the relationship between sleep quality and generalized anxiety symptomology. Scientific reports, 14(1), 13575. 10.1038/s41598-024-63939-3

Buysse, D. J., Reynolds, C. F., 3rd, Monk, T. H., Berman, S. R., & Kupfer, D. J. (1989). The Pittsburgh Sleep Quality Index: a new instrument for psychiatric practice and research. Psychiatry research, 28(2), 193–213. 10.1016/0165-1781(89)90047-4

Cabello, R., Salguero, J. M., Fernández-Berrocal, P., & Gross, J. J. (2013). A Spanish adaptation of the Emotion Regulation Questionnaire. European Journal of Psychological Assessment, 29(4), 234–240. 10.1027/1015-5759/a000150

Calhoun, V. D., Wager, T. D., Krishnan, A., Rosch, K. S., Seymour, K. E., Nebel, M. B., Mostofsky, S. H., Nyalakanai, P., & Kiehl, K. (2017). The impact of T1 versus EPI spatial normalization templates for fMRI data analyses. Human brain mapping, 38(11), 5331–5342. 10.1002/hbm.23737

Carver, C. S., & White, T. L. (1994). Behavioral inhibition, behavioral activation, and affective responses to impending reward and punishment: The BIS/BAS Scales. Journal of Personality and Social Psychology, 67(2), 319–333. 10.1037/0022-3514.67.2.319

Chai, X. J., Castañón, A. N., Ongür, D., & Whitfield-Gabrieli, S. (2012). Anticorrelations in resting state networks without global signal regression. NeuroImage, 59(2), 1420–1428. 10.1016/j.neuroimage.2011.08.048

Chai, Y., Gehrman, P., Yu, M., Mao, T., Deng, Y., Rao, J., Shi, H., Quan, P., Xu, J., Zhang, X., Lei, H., Fang, Z., Xu, S., Boland, E., Goldschmied, J. R., Barilla, H., Goel, N., Basner, M., Thase, M. E., Sheline, Y. I., … Rao, H. (2023). Enhanced amygdala-cingulate connectivity associates with better mood in both healthy and depressive individuals after sleep deprivation. Proceedings of the National Academy of Sciences of the United States of America, 120(26), e2214505120. 10.1073/pnas.2214505120

Chen, G., Adleman, N. E., Saad, Z. S., Leibenluft, E., & Cox, R. W. (2014). Applications of multivariate modeling to neuroimaging group analysis: a comprehensive alternative to univariate general linear model. NeuroImage, 99, 571–588. 10.1016/j.neuroimage.2014.06.027

Chen, I. Y., Jarrin, D. C., Ivers, H., & Morin, C. M. (2017). Investigating psychological and physiological responses to the Trier Social Stress Test in young adults with insomnia. Sleep medicine, 40, 11–22. 10.1016/j.sleep.2017.09.011

Chen, Y., Zhang, L., & Yin, H. (2024). Different emotion regulation strategies mediate the relations of corresponding connections within the default-mode network to sleep quality. Brain imaging and behavior, 18(2), 302–314. 10.1007/s11682-023-00828-9

Cox, R. C., & Olatunji, B. O. (2020). Sleep in the anxiety-related disorders: A meta-analysis of subjective and objective research. Sleep medicine reviews, 51, 101282. 10.1016/j.smrv.2020.101282

Cox R. W. (1996). AFNI: software for analysis and visualization of functional magnetic resonance neuroimages. Computers and biomedical research, an international journal, 29(3), 162–173. 10.1006/cbmr.1996.0014

Cullell, N., Cárcel-Márquez, J., Gallego-Fábrega, C., Muiño, E., Llucià-Carol, L., Lledós, M., Amaut, K. E. U., Krupinski, J., & Fernández-Cadenas, I. (2021). Sleep/wake cycle alterations as a cause of neurodegenerative diseases: A Mendelian randomization study. Neurobiology of aging, 106, 320.e1–320.e12. 10.1016/j.neurobiolaging.2021.05.008

Curtis, B. J., Williams, P. G., & Anderson, J. S. (2019). Neural reward processing in self-reported short sleepers: examination of gambling task brain activation in the Human Connectome Project database. Sleep, 42(9), zsz129. 10.1093/sleep/zsz129

Dawel, A., Shou, Y., Gulliver, A., Cherbuin, N., Banfield, M., Murray, K., Calear, A. L., Morse, A. R., Farrer, L. M., & Smithson, M. (2021). Cause or symptom? A longitudinal test of bidirectional relationships between emotion regulation strategies and mental health symptoms. Emotion (Washington, D.C.), 21(7), 1511–1521. 10.1037/emo0001018

Fan, L., Li, H., Zhuo, J., Zhang, Y., Wang, J., Chen, L., Yang, Z., Chu, C., Xie, S., Laird, A. R., Fox, P. T., Eickhoff, S. B., Yu, C., & Jiang, T. (2016). The Human Brainnetome Atlas: A New Brain Atlas Based on Connectional Architecture. Cerebral cortex (New York, N.Y. : 1991), 26(8), 3508–3526. 10.1093/cercor/bhw157

Flandin, G., & Friston, K. J. (2019). Analysis of family-wise error rates in statistical parametric mapping using random field theory. Human brain mapping, 40(7), 2052–2054. 10.1002/hbm.23839

Friston K.J., Ashburner J., Frith C.D., Poline J.B., Heather J.D., & Frackowiak R.S. (1995). Spatial registration and normalization of images. Human brain mapping, 3(3), 165–189. 10.1002/hbm.460030303

Friston, K. J., Williams, S., Howard, R., Frackowiak, R. S., & Turner, R. (1996). Movement-related effects in fMRI time-series. Magnetic resonance in medicine, 35(3), 346–355. 10.1002/mrm.1910350312

Frossard, J., Renaud, O. (2021). Permutation Tests for Regression, ANOVA, and Comparison of Signals: The permuco Package. Journal of Statistical Software, 99(15), 1–32. 10.18637/jss.v099.i15.

Goldin, P. R., McRae, K., Ramel, W., & Gross, J. J. (2008). The neural bases of emotion regulation: reappraisal and suppression of negative emotion. Biological psychiatry, 63(6), 577–586. 10.1016/j.biopsych.2007.05.031

Goldstein, A. N., Greer, S. M., Saletin, J. M., Harvey, A. G., Nitschke, J. B., & Walker, M. P. (2013). Tired and apprehensive: anxiety amplifies the impact of sleep loss on aversive brain anticipation. The Journal of neuroscience : the official journal of the Society for Neuroscience, 33(26), 10607–10615. 10.1523/JNEUROSCI.5578-12.2013

Greer, S. M., Goldstein, A. N., Knutson, B., & Walker, M. P. (2016). A Genetic Polymorphism of the Human Dopamine Transporter Determines the Impact of Sleep Deprivation on Brain Responses to Rewards and Punishments. Journal of cognitive neuroscience, 28(6), 803–810. 10.1162/jocn_a_00939

Gross, J.J., & John, O.P. (2003). Individual differences in two emotion regulation processes: Implications for affect, relationships, and well-being. Journal of Personality and Social Psychology, 85, 348–362. 10.1037/0022-3514.85.2.348

Grumbach, P., Opel, N., Martin, S., Meinert, S., Leehr, E. J., Redlich, R., Enneking, V., Goltermann, J., Baune, B. T., Dannlowski, U., & Repple, J. (2020). Sleep duration is associated with white matter microstructure and cognitive performance in healthy adults. Human brain mapping, 41(15), 4397–4405. 10.1002/hbm.25132

Guerra-Carrillo, B., Mackey, A. P., & Bunge, S. A. (2014). Resting-state fMRI: a window into human brain plasticity. The Neuroscientist : a review journal bringing neurobiology, neurology and psychiatry, 20(5), 522–533. 10.1177/1073858414524442

Gujar, N., Yoo, S. S., Hu, P., & Walker, M. P. (2011). Sleep deprivation amplifies reactivity of brain reward networks, biasing the appraisal of positive emotional experiences. The Journal of neuroscience : the official journal of the Society for Neuroscience, 31(12), 4466–4474. 10.1523/JNEUROSCI.3220-10.2011x

Hallquist, M. N., Hwang, K., & Luna, B. (2013). The nuisance of nuisance regression: spectral misspecification in a common approach to resting-state fMRI preprocessing reintroduces noise and obscures functional connectivity. NeuroImage, 82, 208–225. 10.1016/j.neuroimage.2013.05.116

Hedges, D. W., Chase, M., Farrer, T. J., & Gale, S. D. (2024). Psychiatric Disease as a Potential Risk Factor for Dementia: A Narrative Review. Brain sciences, 14(7), 722. 10.3390/brainsci14070722

Killgore W. D. (2010). Effects of sleep deprivation on cognition. Progress in brain research, 185, 105–129. 10.1016/B978-0-444-53702-7.00007-5

Kim, S. Y., Lee, K. H., Lee, H., Jeon, J. E., Lee, M. H., Lee, J., Oh, S. M., Lee, Y. J., & Kim, S. J. (2022). Negative life stress, sleep disturbance, and depressive symptoms: The moderating role of anterior insula activity in response to sleep-related stimuli. Journal of affective disorders, 299, 553–558. 10.1016/j.jad.2021.12.072

Kriegeskorte, N., Lindquist, M. A., Nichols, T. E., Poldrack, R. A., & Vul, E. (2010). Everything you never wanted to know about circular analysis, but were afraid to ask. Journal of cerebral blood flow and metabolism : official journal of the International Society of Cerebral Blood Flow and Metabolism, 30(9), 1551–1557. 10.1038/jcbfm.2010.86

Liu, C., Lee, S. H., Loewenstein, D. A., Galvin, J. E., Camargo, C. J., & Alperin, N. (2022). Poor sleep accelerates hippocampal and posterior cingulate volume loss in cognitively normal healthy older adults. Journal of sleep research, 31(4), e13538. 10.1111/jsr.13538

Lischke, A., Pahnke, R., Mau-Moeller, A., Jacksteit, R., & Weippert, M. (2020). Sex-Specific Relationships Between Interoceptive Accuracy and Emotion Regulation. Frontiers in behavioral neuroscience, 14, 67. 10.3389/fnbeh.2020.00067

Ma, X., Jiang, G., Tian, J., Liu, M., Fang, J., Xu, Y., & Song, T. (2021). Convergent and divergent functional connectivity alterations of hippocampal subregions between short-term and chronic insomnia disorder. Brain imaging and behavior, 15(2), 986–995. 10.1007/s11682-020-00306-6

Madrid-Valero, J. J., Kirkpatrick, R. M., González-Javier, F., Gregory, A. M., & Ordoñana, J. R. (2023). Sex differences in sleep quality and psychological distress: Insights from a middle-aged twin sample from Spain. Journal of sleep research, 32(2), e13714. 10.1111/jsr.13714

Meyer, N., Lok, R., Schmidt, C., Kyle, S. D., McClung, C. A., Cajochen, C., Scheer, F. A. J. L., Jones, M. W., & Chellappa, S. L. (2024). The sleep-circadian interface: A window into mental disorders. Proceedings of the National Academy of Sciences of the United States of America, 121(9), e2214756121. 10.1073/pnas.2214756121

Minkel, J. D., Banks, S., Htaik, O., Moreta, M. C., Jones, C. W., McGlinchey, E. L., Simpson, N. S., & Dinges, D. F. (2012). Sleep deprivation and stressors: evidence for elevated negative affect in response to mild stressors when sleep deprived. Emotion (Washington, D.C.), 12(5), 1015–1020. 10.1037/a0026871

Mongrain, V., Carrier, J., & Dumont, M. (2006). Difference in sleep regulation between morning and evening circadian types as indexed by antero-posterior analyses of the sleep EEG. The European journal of neuroscience, 23(2), 497–504. 10.1111/j.1460-9568.2005.04561.x

Morawetz, C., Riedel, M. C., Salo, T., Berboth, S., Eickhoff, S. B., Laird, A. R., & Kohn, N. (2020). Multiple large-scale neural networks underlying emotion regulation. Neuroscience and biobehavioral reviews, 116, 382–395. 10.1016/j.neubiorev.2020.07.001

Motomura, Y., Kitamura, S., Oba, K., Terasawa, Y., Enomoto, M., Katayose, Y., Hida, A., Moriguchi, Y., Higuchi, S., & Mishima, K. (2013). Sleep debt elicits negative emotional reaction through diminished amygdala-anterior cingulate functional connectivity. PloS one, 8(2), e56578. 10.1371/journal.pone.0056578

Nieto-Castanon, A. (2020). FMRI minimal preprocessing pipeline. In Handbook of functional connectivity Magnetic Resonance Imaging methods in CONN (pp. 3–16). Hilbert Press.

Oldfield R. C. (1971). The assessment and analysis of handedness: the Edinburgh inventory. Neuropsychologia, 9(1), 97–113. 10.1016/0028-3932(71)90067-4

Owens, J., Adolescent Sleep Working Group, & Committee on Adolescence (2014). Insufficient sleep in adolescents and young adults: an update on causes and consequences. Pediatrics, 134(3), e921–e932. 10.1542/peds.2014-1696

Palmer, C. A., & Alfano, C. A. (2017). Sleep and emotion regulation: An organizing, integrative review. Sleep medicine reviews, 31, 6–16. 10.1016/j.smrv.2015.12.006

Penny W.D., Friston K.J., Ashburner J.T., Kiebel S.J., & Nichols, T. E. (Eds.) (2011). Statistical parametric mapping: the analysis of functional brain images. Elsevier.

Picó-Pérez, M., Fullana, M. A., Albajes-Eizagirre, A., Vega, D., Marco-Pallarés, J., Vilar, A., Chamorro, J., Felmingham, K. L., Harrison, B. J., Radua, J., & Soriano-Mas, C. (2023). Neural predictors of cognitive-behavior therapy outcome in anxiety-related disorders: a meta-analysis of task-based fMRI studies. Psychological medicine, 53(8), 3387–3395. 10.1017/S0033291721005444

Picó-Pérez, M., Radua, J., Steward, T., Menchón, J. M., & Soriano-Mas, C. (2017). Emotion regulation in mood and anxiety disorders: A meta-analysis of fMRI cognitive reappraisal studies. Progress in neuro-psychopharmacology & biological psychiatry, 79(Pt B), 96–104. 10.1016/j.pnpbp.2017.06.001

Pifer, G. C., Ferrara, N. C., & Kwapis, J. L. (2024). Long-lasting effects of disturbing the circadian rhythm or sleep in adolescence. Brain research bulletin, 213, 110978. 10.1016/j.brainresbull.2024.110978

Power, J. D., Mitra, A., Laumann, T. O., Snyder, A. Z., Schlaggar, B. L., & Petersen, S. E. (2014). Methods to detect, characterize, and remove motion artifact in resting state fMRI. NeuroImage, 84, 320–341. 10.1016/j.neuroimage.2013.08.048

Prather, A. A., Bogdan, R., & Hariri, A. R. (2013). Impact of sleep quality on amygdala reactivity, negative affect, and perceived stress. Psychosomatic medicine, 75(4), 350–358. 10.1097/PSY.0b013e31828ef15b

R Core Team. (2022). R: A language and environment for statistical computing. R Foundation for Statistical Computing, Vienna, Austria. URL https://www.R-project.org/.

Reimann, G. M., Küppers, V., Camilleri, J. A., Hoffstaedter, F., Langner, R., Laird, A. R., Fox, P. T., Spiegelhalder, K., Eickhoff, S. B., & Tahmasian, M. (2023). Convergent abnormality in the subgenual anterior cingulate cortex in insomnia disorder: A revisited neuroimaging meta-analysis of 39 studies. Sleep medicine reviews, 71, 101821. 10.1016/j.smrv.2023.101821

Rolls E. T. (2019). The cingulate cortex and limbic systems for emotion, action, and memory. Brain structure & function, 224(9), 3001–3018. 10.1007/s00429-019-01945-2

Scaini, S., Davies, S., De Francesco, S., Pelucchi, A., Rubino, S., & Battaglia, M. (2025). Altered pain perception and nociceptive thresholds in major depression and anxiety disorders: A meta-analysis. Neuroscience and biobehavioral reviews, 169, 106014. 10.1016/j.neubiorev.2025.106014

Shen, Z., Yang, X., She, T., Zhao, G., Dou, Z., Luo, Y., Lin, W., Dang, W., & Yu, S. (2024). Deficits in brain default mode network connectivity mediate the relationship between poor sleep quality and anxiety severity. Sleep, 47(3), zsad296. 10.1093/sleep/zsad296

Sladky, R., Friston, K. J., Tröstl, J., Cunnington, R., Moser, E., & Windischberger, C. (2011). Slice-timing effects and their correction in functional MRI. NeuroImage, 58(2), 588–594. 10.1016/j.neuroimage.2011.06.078

Spielberger, C. D. (1989). State-Trait Anxiety Inventory: Bibliography (2nd ed.). Palo Alto, CA: Consulting Psychologists Press.

Sollenberger, N. A., Sequeira, S., Forbes, E. E., Siegle, G. J., Silk, J. S., Ladouceur, C. D., Ryan, N. D., Dahl, R. E., Mattfeld, A. T., & McMakin, D. L. (2023). More time awake after sleep onset is linked to reduced ventral striatum response to rewards in youth with anxiety. Journal of child psychology and psychiatry, and allied disciplines, 64(1), 83–90. 10.1111/jcpp.13669

Tavernier, R., & Willoughby, T. (2015). A longitudinal examination of the bidirectional association between sleep problems and social ties at university: the mediating role of emotion regulation. Journal of youth and adolescence, 44(2), 317–330. 10.1007/s10964-014-0107-x

Tempesta, D., Salfi, F., De Gennaro, L., & Ferrara, M. (2020). The impact of five nights of sleep restriction on emotional reactivity. Journal of sleep research, 29(5), e13022. 10.1111/jsr.13022

Telzer, E. H., Fuligni, A. J., Lieberman, M. D., & Galván, A. (2013). The effects of poor quality sleep on brain function and risk taking in adolescence. NeuroImage, 71, 275–283. 10.1016/j.neuroimage.2013.01.025

Tiede, W., Magerl, W., Baumgärtner, U., Durrer, B., Ehlert, U., & Treede, R. D. (2010). Sleep restriction attenuates amplitudes and attentional modulation of pain-related evoked potentials, but augments pain ratings in healthy volunteers. Pain, 148(1), 36–42. 10.1016/j.pain.2009.08.029

Tomaso, C. C., Johnson, A. B., & Nelson, T. D. (2021). The effect of sleep deprivation and restriction on mood, emotion, and emotion regulation: three meta-analyses in one. Sleep, 44(6), zsaa289. 10.1093/sleep/zsaa289

Torrubia, R., Ávila, C., Moltó, J. & Caseras, X. (2001). The Sensitivity to Punishment and Sensitivity to Reward Questionnaire (SPSRQ) as a measure of Gray’s anxiety and impulsivity dimensions. Personality and Individual Differences, 31, 837–862. 10.1016/S0191-8869(00)00183-5

Tyra, A. T., Fergus, T. A., & Ginty, A. T. (2024). Emotion suppression and acute physiological responses to stress in healthy populations: a quantitative review of experimental and correlational investigations. Health psychology review, 18(2), 396–420. 10.1080/17437199.2023.2251559

Uddin, L. Q., Yeo, B. T. T., & Spreng, R. N. (2019). Towards a Universal Taxonomy of Macro-scale Functional Human Brain Networks. Brain topography, 32(6), 926–942. 10.1007/s10548-019-00744-6

Vargas, I., & Lopez-Duran, N. (2017). Investigating the effect of acute sleep deprivation on hypothalamic-pituitary-adrenal-axis response to a psychosocial stressor. Psychoneuroendocrinology, 79, 1–8. 10.1016/j.psyneuen.2017.01.030

Vazsonyi, A. T., Liu, D., & Blatny, M. (2022). Longitudinal bidirectional effects between sleep quality and internalizing problems. Journal of adolescence, 94(3), 448–461. 10.1002/jad.12039

Venkatraman, V., Chuah, Y. M., Huettel, S. A., & Chee, M. W. (2007). Sleep deprivation elevates expectation of gains and attenuates response to losses following risky decisions. Sleep, 30(5), 603–609. 10.1093/sleep/30.5.603

Venkatraman, V., Huettel, S. A., Chuah, L. Y., Payne, J. W., & Chee, M. W. (2011). Sleep deprivation biases the neural mechanisms underlying economic preferences. The Journal of neuroscience : the official journal of the Society for Neuroscience, 31(10), 3712–3718. 10.1523/JNEUROSCI.4407-10.2011

Vulser, H., Lemaître, H. S., Guldner, S., Bezivin-Frère, P., Löffler, M., Sarvasmaa, A. S., Massicotte-Marquez, J., Artiges, E., Paillère Martinot, M. L., Filippi, I., Miranda, R., Stringaris, A., van Noort, B. M., Penttilä, J., Grimmer, Y., Becker, A., Banaschewski, T., Bokde, A. L. W., Desrivières, S., Fröhner, J. H., … IMAGEN Consortium (2023). Chronotype, Longitudinal Volumetric Brain Variations Throughout Adolescence, and Depressive Symptom Development. Journal of the American Academy of Child and Adolescent Psychiatry, 62(1), 48–58. 10.1016/j.jaac.2022.06.003

Wang, W., Zhornitsky, S., Li, C. S., Le, T. M., Joormann, J., & Li, C. R. (2019). Social anxiety, posterior insula activation, and autonomic response during self-initiated action in a Cyberball game. Journal of affective disorders, 255, 158–167. 10.1016/j.jad.2019.05.046

Wang, Y., Jiang, P., Tang, S., Lu, L., Bu, X., Zhang, L., Gao, Y., Li, H., Hu, X., Wang, S., Jia, Z., Roberts, N., Huang, X., & Gong, Q. (2021). Left superior temporal sulcus morphometry mediates the impact of anxiety and depressive symptoms on sleep quality in healthy adults. Social cognitive and affective neuroscience, 16(5), 492–501. 10.1093/scan/nsab012

Whitfield-Gabrieli, S., & Nieto-Castanon, A. (2012). Conn: a functional connectivity toolbox for correlated and anticorrelated brain networks. Brain connectivity, 2(3), 125–141. 10.1089/brain.2012.0073

Whitfield-Gabrieli, S., Nieto-Castanon, A., & Ghosh, S. (2011). Artifact detection tools (ART). Cambridge, MA. Release Version, 7(19), 11.

Wright, C. E., Valdimarsdottir, H. B., Erblich, J., & Bovbjerg, D. H. (2007). Poor sleep the night before an experimental stress task is associated with reduced cortisol reactivity in healthy women. Biological psychology, 74(3), 319–327. 10.1016/j.biopsycho.2006.08.003

Wu, Y., Zhuang, Y., & Qi, J. (2020). Explore structural and functional brain changes in insomnia disorder: A PRISMA-compliant whole brain ALE meta-analysis for multimodal MRI. Medicine, 99(14), e19151. 10.1097/MD.0000000000019151

Yarkoni, T., Poldrack, R. A., Nichols, T. E., Van Essen, D. C., & Wager, T. D. (2011). Large-scale automated synthesis of human functional neuroimaging data. Nature methods, 8(8), 665–670. 10.1038/nmeth.1635

Yin, X., Jiang, T., Song, Z., Zhu, L., Wang, G., & Guo, J. (2024). Increased functional connectivity within the salience network in patients with insomnia. Sleep & breathing = Schlaf & Atmung, 28(3), 1261–1271. 10.1007/s11325-024-03002-7

Yoo, S. S., Gujar, N., Hu, P., Jolesz, F. A., & Walker, M. P. (2007). The human emotional brain without sleep--a prefrontal amygdala disconnect. Current biology : CB, 17(20), R877–R878. 10.1016/j.cub.2007.08.007

Zareba, M. R., Fafrowicz, M., Marek, T., Beldzik, E., Oginska, H., & Domagalik, A. (2022). Late chronotype is linked to greater cortical thickness in the left fusiform and entorhinal gyri. Biological Rhythm Research, 53(10), 1626–1638. 10.1080/09291016.2021.1990501

Zareba, M. R., Bielski, K., Costumero, V., & Visser, M. (2024a). Graph analysis of guilt processing network highlights links with subclinical anxiety and self-blame. Social cognitive and affective neuroscience, 19(1), nsae092. 10.1093/scan/nsae092

Zareba, M. R., Fafrowicz, M., Marek, T., Oginska, H., Beldzik, E., & Domagalik, A. (2024b). Tracing diurnal differences in brain anatomy with voxel-based morphometry - associations with sleep characteristics. Chronobiology international, 41(2), 201–212. 10.1080/07420528.2024.2301944

Zhang, L., Li, D., & Yin, H. (2020). How is psychological stress linked to sleep quality? The mediating role of functional connectivity between the sensory/somatomotor network and the cingulo-opercular control network. Brain and cognition, 146, 105641. 10.1016/j.bandc.2020.105641

Zhang, L., Cao, G., Liu, Z., Bai, Y., Li, D., Liu, J., & Yin, H. (2022). The gray matter volume of bilateral inferior temporal gyrus in mediating the association between psychological stress and sleep quality among Chinese college students. Brain imaging and behavior, 16(2), 557–564. 10.1007/s11682-021-00524-6

