## Supplementary Material for "Subjective sleep quality in healthy young adults moderates associations of sensitivity to punishment and reward with functional connectivity of regions relevant for insomnia disorder"

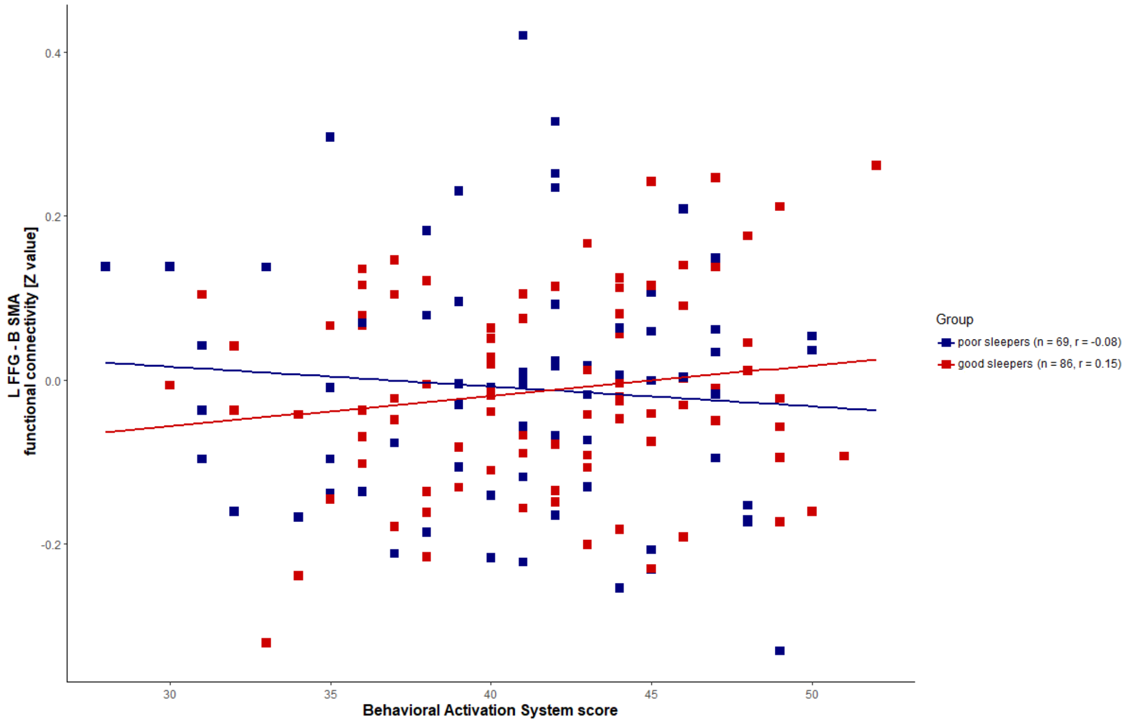


**Supplementary Figure 1.** The effects of interactions between subjective sleep quality and positive reinforcement processing, as measured with the Behavioral Activation System scale (Carver and White, 1994), on the functional connectivity between the left fusiform gyrus (L FFG) and bilateral supplementary motor area (B SMA). While in the individuals with good sleep quality the association was positive, its direction was reversed in the poor sleepers, therefore replicating the effects observed for sensitivity to reward in the primary analysis ($F = 7.53$; $p_{FDR} = 0.03$; $\eta p^2 = 0.049$).

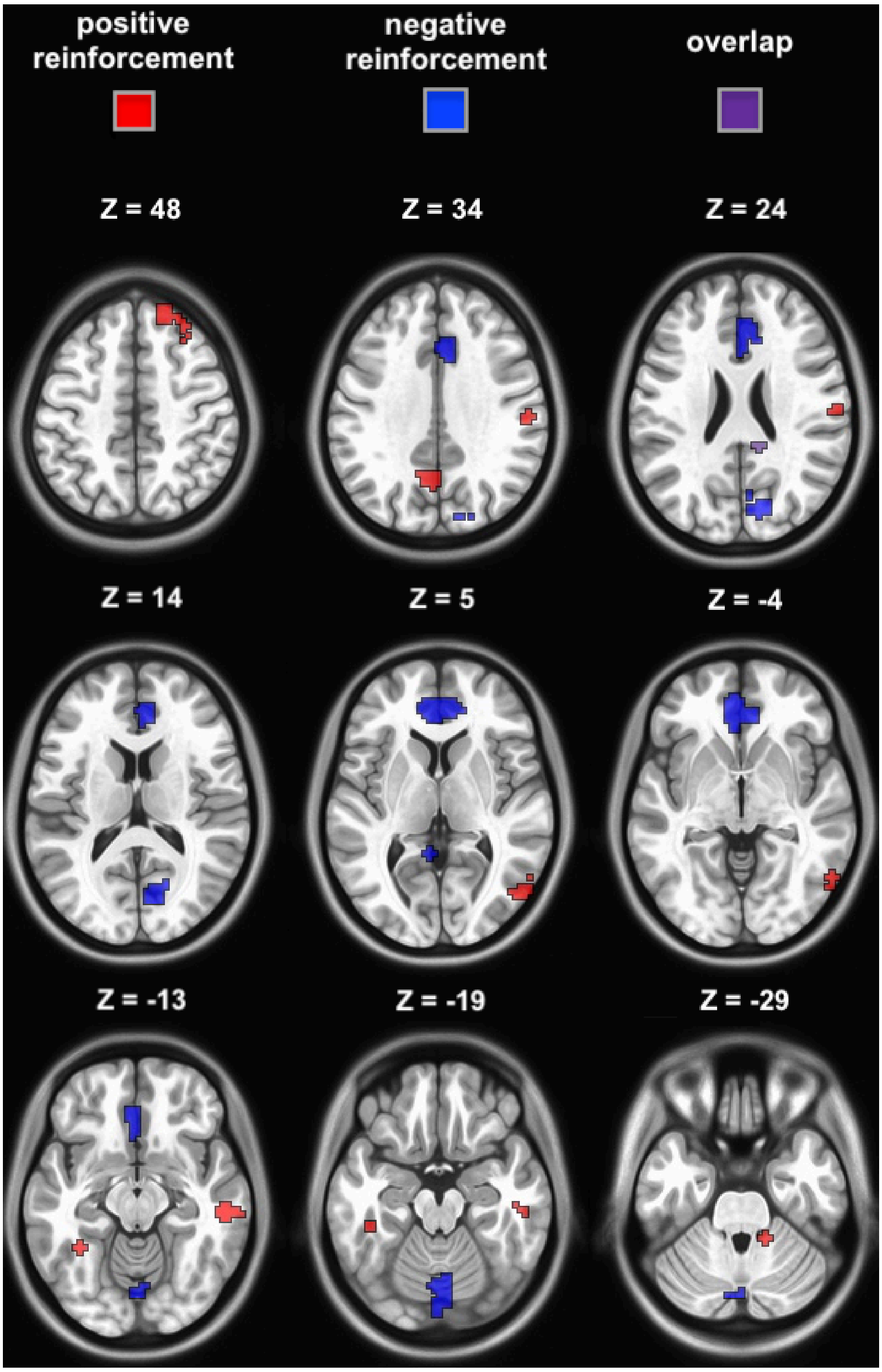


**Supplementary Figure 2.** Brain areas with their functional connectivity patterns significantly associated with the positive and negative reinforcement processing. The figure shows a combination of seed masks and significant results for the following regions of interest: bilateral subgenual anterior and posterior cingulate cortex, right pregenual anterior cingulate cortex, middle temporal gyrus and anterior lobe of cerebellum, and the left fusiform gyrus.

**Supplementary Table 1.** The results of the post-hoc analysis for the two-way interactions of reinforcement sensitivity and the sleep quality group. Slopes of the associations between the reward or punishment sensitivity and functional connectivity were compared between the good and poor sleepers.

| **Functional connectivity pattern** | **Reinforcement type** | **Good sleepers**<br><br>**β (SE) [95% CI]** | **Poor sleepers**<br><br>**β (SE) [95% CI]** | **T-ratio (p value)** |
|---|---|---|---|---|
| L fusiform gyrus BA 20<br>B supplementary motor area | Reward | 0.009 (0.003)<br>[0.002; 0.016] | -0.017 (0.004)<br>[-0.024; -0.009] | 4.97 (< 0.0001) |
| L precentral gyrus<br>L supramarginal gyrus, L superior temporal gyrus | Punishment | 0.012 (0.003)<br>[0.006; 0.018] | -0.013 (0.004)<br>[-0.020; -0.005] | 5.09 (< 0.0001) |
| L precentral gyrus<br>L posterior insula | Punishment | 0.006 (0.002)<br>[0.001; 0.011] | -0.013 (0.003)<br>[-0.019; -0.008]] | 5.51 (< 0.0001) |

Abbreviations: SE; standard error; CI, confidence intervals; R, right; L, left; B, bilateral; BA, Brodmann area.

**Supplementary Table 2.** The main effects of positive and negative reinforcement processing on the functional connectivity of regions of interests. The primary whole-brain models were tested using the Sensitivity to Reward and Sensitivity to Punishment scales (Torrubia et al., 2001). The replication analyses were run with the Behavioral Activation and Behavioral Inhibition System scores (Carver and White, 1994) using the *aovperm* function from the permuco library (Frossard and Renaud, 2021) in R (version 4.2.1; R Core Team, 2022) with 10000 permutations. The replication models included only the main effects of the reinforcement processing measures, i.e. the interaction terms were not tested. Raw, uncorrected p values are provided. Asterisks indicate significance after false discovery rate (FDR < 0.05) correction.

| **Region of interest** | **Reinforcement type** | **MNI** | **Location** | **Voxels** | **Whole-brain model F-stat (Pearson r)** | **Replication model F-stat** | **Replication model p value** | **Replication model ηp²** |
|---|---|---|---|---|---|---|---|---|
| R posterior cingulate cortex | positive | 52, -72, 6 | R lateral occipital cortex | 51 | 18.84 (0.386) | 8.99 | 0.003* | 0.057 |
| R middle temporal gyrus | positive | 22, 44, 48 | R superior frontal gyrus | 49 | 25.05 (0.394) | 2.67 | 0.105 | 0.018 |
| R anterior lobe of cerebellum | positive | 54, -24, -12 | R middle temporal gyrus | 40 | 26.42 (0.383) | 9.15 | 0.003* | 0.058 |
| L fusiform gyrus BA 37 | positive | 58, -22, 36 | R supramarginal gyrus | 42 | 27.69 (0.377) | 9.92 | 0.002* | 0.062 |
| L fusiform gyrus BA 20 | positive | -6, -64, 32 | L precuneus | 39 | 20.13 (0.319) | 1.32 | 0.247 | 0.009 |
| L posterior cingulate cortex | negative | 12, -78, 18 | R cuneus | 66 | 22.94 (-0.361) | 5.19 | 0.022* | 0.034 |
| L subgenual anterior cingulate cortex | negative | -2, -72, -18 | B cerebellar lobule VI (vermis) | 55 | 20.78 (-0.487) | 10.93 | 0.001* | 0.068 |
| R subgenual anterior cingulate cortex | negative | 0, -76, -16 | B cerebellar lobule VI (vermis) | 52 | 23.13 (-0.467) | 5.61 | 0.017* | 0.036 |
| R pregenual anterior cingulate cortex | negative | 0, -82, -18 | B cerebellar lobule VI (vermis) | 44 | 20.79 (-0.388) | 5.81 | 0.016* | 0.038 |
| R posterior cingulate cortex | negative | 12, -82, 26 | R cuneus | 78 | 23.76 (-0.354) | 6.91 | 0.009* | 0.044 |

Abbreviations: R, right; L, left; B, bilateral; BA, Brodmann area.

**Supplementary Table 3.** The associations of trait anxiety, and the use of emotion reappraisal and suppression strategies with the functional connectivity patterns linked to sleep quality and its interactions with reinforcement processing measures. False discovery rate (FDR)-corrected p values are provided.

| **Functional connectivity pattern** | **Factor** | **Trait anxiety T-stat (FDR)** | **Emotion reappraisal T-stat (FDR)** | **Emotion suppression T-stat (FDR)** |
|---|---|---|---|---|
| R subgenual anterior cingulate cortex<br>L prefrontal thalamus | Sleep quality | -0.65 (0.646) | 0.56 (0.719) | -3.22 (0.01) |
| L fusiform gyrus BA 20<br>B supplementary motor area | Sleep quality × SR | -1.61 (0.278) | 1.64 (0.358) | 1.22 (0.375) |
| L precentral gyrus<br>L supramarginal gyrus, L superior temporal gyrus | Sleep quality × SP | -1.37 (0.288) | 1.47 (0.358) | 1.29 (0.375) |
| L precentral gyrus<br>L posterior insula | Sleep quality × SP | -2.76 (0.033) | 0.83 (0.685) | -0.61 (0.544) |
| R middle temporal gyrus<br>L lateral prefrontal cortex | Sleep quality × SP × SR | 0.35 (0.728) | -0.14 (0.892) | -0.81 (0.526) |

Abbreviations: R, right; L, left; B, bilateral; BA, Brodmann area; SR, sensitivity to reward; SP, sensitivity to punishment.

**Supplementary Table 4.** The associations of the successfully replicated positive and negative reinforcement processing-related functional connectivity patterns with trait anxiety, and the use of emotion reappraisal and suppression strategies. Raw, uncorrected p values are provided.

| **Functional connectivity pattern** | **Reinforcement type** | **Trait anxiety T-stat (p value)** | **Emotion reappraisal T-stat (p value)** | **Emotion suppression T-stat (p value)** |
|---|---|---|---|---|
| R posterior cingulate cortex<br>R lateral occipital cortex | positive | -0.41 (0.681) | -0.43 (0.67) | -0.08 (0.935) |
| R anterior lobe of cerebellum<br>R middle temporal gyrus | positive | -0.83 (0.407) | 0.70 (0.488) | -0.61 (0.545) |
| L fusiform gyrus BA 37<br>R supramarginal gyrus | positive | 1.98 (0.049) | -0.39 (0.7) | -2.26 (0.025) |
| L posterior cingulate cortex<br>R cuneus | negative | -2.46 (0.015) | 0.48 (0.634) | -2.9 (0.004) |
| L subgenual anterior cingulate cortex<br>B cerebellar lobule VI (vermis) | negative | -6.47 (<0.0001) | 0.84 (0.401) | -3.43 (0.0007) |
| R subgenual anterior cingulate cortex<br>B cerebellar lobule VI (vermis) | negative | -5.00 (<0.0001) | -0.02 (0.985) | -3.53 (0.0006) |
| R pregenual anterior cingulate cortex<br>B cerebellar lobule VI (vermis) | negative | -3.89 (0.0001) | -0.70 (0.484) | -2.68 (0.008) |
| R posterior cingulate cortex<br>R cuneus | negative | -2.97 (0.003) | 1.08 (0.281) | -1.39 (0.166) |

Abbreviations: R, right; L, left; B, bilateral; BA, Brodmann area.
